# Fast Targeted Metabolomics for Analyzing Metabolic Diversity of Bacterial Indole Derivatives in ME/CFS Gut Microbiome

**DOI:** 10.1101/2024.07.29.605643

**Authors:** Huidi Tian, Lei Wang, Elizabeth Aiken, Robert Jervine V. Ortega, Rachel Hardy, Lindsey Placek, Lina Kozhaya, Derya Unutmaz, Julia Oh, Xudong Yao

## Abstract

Disruptions in microbial metabolite interactions due to gut microbiome dysbiosis and metabolomic shifts may contribute to Myalgic Encephalomyelitis/Chronic Fatigue Syndrome (ME/CFS) and other immune-related conditions. The aryl hydrocarbon receptor (AhR), activated upon binding various tryptophan metabolites, modulates host immune responses. This study investigates whether the metabolic diversity—the concentration distribution—of bacterial indole pathway metabolites can differentiate bacterial strains and classify ME/CFS samples. A fast targeted liquid chromatography-parallel reaction monitoring method at a rate of 4 minutes per sample was developed for large-scale analysis. This method revealed significant metabolic differences in indole derivatives among *B. uniformis* strains cultured from human isolates. Principal component analysis identified two major components (PC1, 68.9%; PC2, 18.7%), accounting for 87.6% of the variance and distinguishing two distinct *B. uniformis* clusters. The metabolic difference between clusters was particularly evident in the relative contributions of indole-3-acrylate and indole-3-aldehyde. We further measured concentration distributions of indole derivatives in ME/CFS by analyzing fecal samples from 10 patients and 10 healthy controls using the fast targeted metabolomics method. An AdaBoost-LOOCV model achieved moderate classification success with a mean LOOCV accuracy of 0.65 (Control: precision of 0.67, recall of 0.60, F1-score of 0.63; ME/CFS: precision of 0.64, recall of 0.7000, F1-score of 0.67). These results suggest that the metabolic diversity of indole derivatives from tryptophan degradation, facilitated by the fast targeted metabolomics and machine learning, is a potential biomarker for differentiating bacterial strains and classifying ME/CFS samples. Mass spectrometry datasets are accessible at the National Metabolomics Data Repository (ST002308, DOI: 10.21228/M8G13Q; ST003344, DOI: 10.21228/M8RJ9N; ST003346, DOI: 10.21228/M8RJ9N).

## INTRODUCTION

Gut microbiome dysbiosis and shifts, associated with metabolomic changes, manifest in a range of infectious and chronic diseases,^1–4^ including poorly understood, chronic debilitating conditions such as Myalgic Encephalomyelitis/Chronic Fatigue Syndrome (ME/CFS).^5–8^ Disruptions in microbial metabolite interactions with the aryl hydrocarbon receptor (AhR) are posited to contribute to ME/CFS and other immune-related diseases like cancer;^9,10^ thus AhR serves as a therapeutic target.^11–16^ AhR is a transcriptional factor that activates upon ligand binding, leading to the modulation of the host immune response.^17–19^ It binds a diverse array of ligands, including many tryptophan metabolites such as kynurenine, 5-hydroxytryptophan, indole-3-acetate, and indole-3-propionate.

Tryptophan is mainly metabolized via three pathways: serotonin, kynurenine, and indole.^20^ While host-microbe co-metabolism heavily affects the serotonin and kynurenine pathways, bacterial indole derivatives provide a more defined molecule set for elucidating microbial impact on host health. Recent reports on targeted quantitation of tryptophan metabolites in various biomatrices typically employ liquid chromatography-selected reaction monitoring mass spectrometry (LC-SRM MS) on triple quadrupole mass spectrometers. This approach has proven utility in clinical settings, with the analysis rate ranging from 7 to 15 minutes per sample.^21–25^ For instance, methods with a 7-minute run time allow the analysis of two 96-well plates in 24 hours, quantifying 18 tryptophan metabolites for a clinical study of inflammatory bowel disease,^23^ while another 7-minute method targets 31 metabolites in the tryptophan pathways.^22^ A longer 15-minute method enables the analysis of 89 target metabolites, though at the expense of sample throughput.^25^

High-resolution, high-accuracy mass spectrometers offer alternative methods for targeted quantitation of tryptophan metabolites while still providing sufficient limits of quantitation. Unlike SRM using unit-resolution quadrupole analyzers, which rely on selective transitions for selectivity, high-resolution methods achieve selectivity based on the mass accuracy and resolution of fragment ions. The added separation capability from the mass-to-charge dimension better tolerates the selectivity pressure resulting from reduced chromatographic separation time for increased sample throughput. Additionally, these methods facilitate post-acquisition spectral analysis and allow for tuning of method selectivity via fragment ion usage. Methods using quadrupole-time of flight (QTOF)^26^ and quadrupole-orbitrap^27^ instruments have been reported. A liquid chromatography-parallel reaction monitoring (LC-PRM) mass spectrometry method with a 15-minute run time quantifies neurotransmitters and tryptophan metabolites in mouse serum, feces, and brain,^27^ providing insights to the microbiota-gut-brain axis.^28^

This study investigates whether the local metabolic diversity—defined as the percentage concentration distribution of metabolites—in the bacterial indole pathway can be used to differentiate bacterial strains and classify fecal samples from ME/CFS patients versus healthy controls. Our assumptions are that (1) the genetic regulation of tryptophan degradation is distinctive to bacteria, and (2) the synergistic interplay of indole derivatives from gut microbiome significantly influences the regulation of AhR in the human host.

Utilizing the metabolic diversity of indole derivatives necessitates accurately and precisely measuring the relative concentrations of these compounds in complex biomatrices, as well as conducting high-throughput analysis on a large scale to obtain biologically significant data. In this work, LC-PRM-based targeted quantitation of indole derivatives from the tryptophan indole pathway—reflecting the local metabolic diversity of bacterial indole derivatives—is utilized to establish a high-resolution method for differentiating bacterial strains. Sample throughput is further enhanced by developing a fast targeted PRM method at a throughput rate of 4 minutes per sample, aiming at large-scale analysis. Overall, fast targeted metabolomics of indole derivatives in patient and control fecal samples, assisted by machine learning, reveals the local metabolic diversity of indole derivatives as a potential biomarker for ME/CFS.

## EXPERIMENTAL SECTION

### Bacterial Culture and Supernatant Preparation

Bacterial strains were cultured from the microbiota of healthy volunteers or ME/CFS patients at Jackson Laboratory for Genomic Medicine (Farmington, CT). The derived bacteria were cultured overnight in tryptic soy broth (TSB) media or TSB media supplemented with vitamin K (0.1 mg/L) and hemin (5 mg/L) to saturation under anaerobic or aerobic conditions at 37℃, as appropriate. OD600 was used to estimate cell density, and 1 mL of cells were pelleted. The supernatant was filtered through a 0.22-µm filter to prepare cell-free culture supernatants, which were then stored at -80℃ . Preparation of full extracts and HPLC fractions followed our previous publication.^29^

### Fecal Sample Extraction

Fecal samples from 10 healthy cohorts and 10 ME/CFS patients were lyophilized overnight. Sixty 1.5 mL microcentrifuge tubes were weighed and labeled. Each sample was homogenized with an ultrasonication probe and approximately 20 mg was aliquoted into a microcentrifuge tube in triplicate. The tubes were weighed again to obtain the exact mass. Ice-cold 50% (v/v) methanol in water was added to each tube in a ratio of 600 µL extraction solvent per 25 mg fecal sample. Approximately 10 2-mm zirconium beads were added to each tube, and the samples were extracted with a homogenizer for 10 minutes at 30 Hz. The resulting solutions were centrifuged at 12,000 x g for 10 minutes. The supernatants were filtered with a 0.22 µm filter. The resulting solution was dried by Speedvac and reconstituted in 2% acetonitrile (MeCN) at a concentration factor of 5 for further analysis.

### Solid Phase Extraction

Solid phase extraction (SPE) was performed to prepare cell-free culture supernatants for LC-MS analyses. Reversed-phase SPE was conducted using Targa C18 sorbent (Higgins Analytical, The Nest Group). Each commercial Targa C18 reversed-phase MacroSpin column was packed with 200 mg of Targa C18 sorbent, with a bed volume of 200 µL. The sorbent was conditioned with 200 µL of methanol three times, followed by centrifugation at 110 × g for 2 minutes. This was followed by equilibration with 200 µL of water, repeated thrice (2 minutes at 110 × g). Samples were loaded into each prepared column and centrifuged at 110 × g for 2 minutes. After loading, each cartridge was washed with 200 µL of water. The sample was eluted with 200 µL of 80% MeCN in water (v/v), and the eluates were collected (110 × g for 2 minutes). The eluates were dried using a SpeedVac and lyophilizer for future analysis.

### Solid Phase Extraction in Plate Format

High-throughput reversed-phase SPE was performed using a Targa C18 (Higgins Analytical, The Nest Group) 96-well plate format. Each well was conditioned with 200 µL of methanol three times with centrifugation at 600 × g for 2 minutes, followed by equilibration with 200 µL of water (2 minutes at 800 × g) three times. The samples were loaded into each prepared well with centrifugation at 800 × g for 2 minutes. After loading, each well was washed with 200 µL of water five times. The sample was eluted with 200 µL of 80% MeCN in water (v/v), and the eluates were collected in a 96-deep well plate (800 × g for 2 minutes). The resulting solution was dried and reconstituted with 2% MeCN for further analysis.

### Synthesis of Isotopically Labeled N-Formylkynurenine (NFK-C13)

The synthesis of isotopically labeled NFK-C13 and its detailed characterization were previously described.^29^ Briefly, 100 µL of 20 mM KYN in 2-(N-morpholino)ethanesulfonic acid (MES) buffer (1 M, pH 6.1) were mixed with 50 µL of formic acid-C13. The reaction mixture was incubated in a thermomixer at 800 rpm and room temperature overnight. The solution was then dried by overnight lyophilization, and the resulting solid was reconstituted with water for analysis.

### Metabolomic Profiling using Data-Dependent Acquisition of Bacteria Cultural Supernatants and Fractions

The bacterial culture supernatant full extracts and extract fractions of *Staphylococcus epidermidis* (*S. epidermidis*) and *Enterococcus faecium* (*E. faecium*), isolated from a human host and cultured, were prepared in a previous study.^29^ The culture supernatant of *S. epidermidis* had relatively high AhR-activating potency, and *E. faecium* had low potency. The full extracts and fractionated samples were analyzed on a quadrupole-orbitrap mass spectrometer (Exploris 480, Thermo Fisher Scientific) equipped with the heated electrospray ionization (H-ESI) source and coupled to a Vanquish UPLC system (Thermo Fisher). A reversed-phase column (CORTECS UPLC T3, 2.1 x 150 mm, 1.6 μm particle size, 0.52 mL of bed volume, Waters) was used. The autosampler temperature was 4.0 °C, and the column oven temperature was 40.0 °C. The sample injection volume was 10 µL. The mobile phase flow rate was 300 µL/min. Solvent A was 0.1% formic acid (FA) in water, and solvent B was 0.1% FA in MeCN. The gradient (% for Solvent B at runtime) method was 1% from 0 to 1 minute, 20% at 15 minutes, 90% from 15.1 to 19 minutes, and 1% from 19.1 to 21 minutes.

MS and data-dependent acquisition (DDA) MS/MS modes were used to analyze the full extracts and fractions in both positive and negative ionization modes. The spray voltage was set to static, with 3500 V for the positive mode and 2500 V for the negative mode. The sheath gas was set to 50 arb, the aux gas was set to 10 arb, and the sweep gas was set to 1 arb. The ion transfer tube temperature was set to 300 ℃, and the vaporizer temperature was set to 350 ℃. In the full MS mode, the orbitrap resolution was set to 120,000, and the scan range was 100-1000 m/z. The RF lens was set to 50%, and the AGC target was set to standard. The maximum injection time mode was set to auto. The microscan was set to 1. The data was collected in profile format. In the DDA mode, a 0.7 m/z isolation window was used without isolation offset. The collision energy was set as a stepped mode with normalized HCD collision energy, 15%, 30%, and 45%. The orbitrap resolution was set to 15,000, and the scan range mode was auto-controlled. The AGC target was set as standard, and the maximum injection time mode was set to auto. The intensity threshold was set to 1e4, and the dynamic exclusion was set to 6 seconds. The microscan was set to 1. The data was collected in profile mode.

Data processing was performed using Compound Discoverer software (Version 3.3.3.200, Thermo Fisher Scientific). Raw data files obtained in both polarities for each extract (full and fractionated) were processed simultaneously (mixed polarity). Filters used for metabolite intensity comparison: z = +1 only; ppm error tolerance = ± 5 ppm; retention time between 0.9 - 16.50 mins; peaks not found in blank; peaks should be found in at least 2 replicates with a peak score greater than or equal to 5. Metabolika pathways were used to plot the detected compounds into the relevant pathways. The pathways of interest were the built-in "superpathway for indole-3-acetate conjugate biosynthesis" and an imported bacterial tryptophan catabolism pathway.^20^ Results from the volcano plot of the differential analysis of *S. epidermidis* versus *E. faecium* were replotted using RStudio (Version 2023.06.0, Posit Software, PBC).

### Standard 28-Minute LC-Parallel Reaction Monitoring Method for Targeted Quantitation for Tryptophan Metabolite Quantitation and Comparing Metabolic Diversity among Genera, Species, and Strains

Targeted quantitation of indole compounds in full extracts of culture supernatants across various genera, species, and strains (**Table S1**) referenced a method of LC-PRM MS.^27^ The supernatant samples were analyzed in triplicate either as 5-fold diluted or 5-fold concentrated compared to the initial supernatants. The spiked N-formylkynurenine-C13 (NFK-C13) as the quantitation reference was 0.50 µM in each sample. An equal molarity (10 µM) stock solution of metabolite reference compounds was prepared. Twelve dilutions of this stock, each spiked with the same amount of NFK-C13 (0.50 µM), were measured to generate calibration curves. The calibration curves for each metabolite contained at least five concentration levels (**Figure S1**, **Table S2**).

Targeted quantitation of tryptophan compounds in the PRM mode used a quadrupole-orbitrap mass spectrometer (Exploris 480, Thermo Fisher Scientific) with an H-ESI ionization source in the positive ion mode. HPLC used a reversed-phase column (CORTECS UPLC T3, 2.1 x 150 mm, 1.6 μm particle size, 0.52 mL of bed volume, Waters). The autosampler temperature was 4.0 °C, and the column oven temperature was 40.0 °C. The sample injection volume was 10 µL. The mobile phase flow rate was 300 µL/min. Solvent A was 0.1 % FA in water, and solvent B was 0.1 % FA in MeCN. The gradient (% for Solvent B at runtime) method was 1% from -8 to 0 minute for the equilibration (8 minutes before sample injection), 1% from 0 to 1 minute, 90% at 15 minutes, 90% from 15.1 to 19 minutes, and 1% from 19.1 to 20 minutes. Major mass spectrometer parameters were 3500 V for the spray voltage, 50% for RF lens, 50 arb for the sheath gas, 10 arb for the aux gas, 1 arb for the sweep gas, 300 ℃ for the ion transfer tube, and 350 ℃ for the vaporizer temperature. The isolation window was set to 0.7 m/z without isolation offset. The collision energy was set to fix; the normalized collision energy was optimized for each compound and was shown in **Table S2**. The orbitrap resolution was 15,000, and the scan range mode was auto. The AGC target was set as standard, the maximum injection time mode was set to auto, the intensity threshold was 1e^4^, the dynamic exclusion was 6 seconds, and the microscan was set to 1. The data was collected in profile mode. Quantitation used Skyline.^30^ The data is available at the Metabolomics Workbench.^31^ Study ID: ST002308; doi: 10.21228/M8G13Q.

### Fast 4-Minute LC-Parallel Reaction Monitoring Method for Targeted Quantitation of Indole Compounds in Full Extracts of *Bacteroides uniformis* and Fecal Samples

Full extracts of culture supernatants from various *Bacteroides uniformis* (*B. uniformis*) strains or fecal samples were spiked with NFK-C13, indole-3-propoinic acid-2D (IPA-2D), and indole-3-carboxaldehyde-8C13 (ICA-8C13), using an Opentron OT2 robot and desalted using Targa C18 plates. The desalted extracts were dried by lyophilization and reconstituted with 2% MeCN for LC-PRM MS analysis of a panel of indole compounds (**Table 1**). The supernatants were analyzed at a concentration factor. The concentration of each spiked standard was 0.50 µM per sample. Targeted quantitation of selected indole compounds was performed on a quadrupole-orbitrap mass spectrometer (Exploris 480, Thermo Fisher Scientific) with an H-ESI ionization source in positive ion mode. The front-end separation utilized a reversed-phase column (CORTECS UPLC T3, 2.1 x 50 mm, 1.6 μm particle size, Waters). The autosampler temperature was maintained at 4.0 °C, and the column oven temperature was set at 40.0 °C. The sample injection volume was 5 µL. The mobile phase flow rate was 1 mL/min. Solvent A was 0.1% FA in water, and solvent B was 0.1% FA in MeCN. The gradient (% for Solvent B at runtime) was as follows: 1% from -1.5 to 0 minutes for equilibration, 1% from 0 to 0.2 minutes, 65% at 1.7 minutes, 90% from 1.8 to 2.2 minutes, and 1% from 2.3 to 2.5 minutes.

**Table 1.** Selected indole compounds for fast targeted quantitation.

| Compound Name | Theoretical *m/z* | Linear Range (nM) | R<sup>2</sup> | Normalized CE (%) | RT (min) | RT Window (min) |
| --- | --- | --- | --- | --- | --- | --- |
| N-formylkynurenine-C13 | 238.0909 | - | - | 30 | 0.56 | 0.30 |
| N-formylkynurenine | 237.0870 | 2.5 - 10000 | 0.9981 | 30 | 0.78 | 0.30 |
| Kynurenine | 209.0926 | 2.5 - 10000 | 0.9966 | 30 | 0.78 | 0.30 |
| Indole-3-acetonitrile | 157.0766 | 500 - 10000 | 0.9996 | 80 | 1.55 | 0.15 |
| Indole-3-acrylic acid | 188.0712 | 5 - 10000 | 0.9932 | 50 | 1.49 | 0.20 |
| Indole-3-ethanol | 162.0919 | 5 - 10000 | 0.9986 | 40 | 1.42 | 0.15 |
| Indole-3-propionic acid | 190.0868 | 2.5 - 10000 | 0.9955 | 60 | 1.54 | 0.15 |
| Indole-3-propionic acid-2D | 192.0994 | - | - | 60 | 1.54 | 0.15 |
| Indole-3-acetic acid | 176.0712 | 2.5 - 10000 | 0.9937 | 90 | 1.42 | 0.20 |
| Indole-3-butyric acid | 204.1025 | 10 - 10000 | 0.9957 | 80 | 1.61 | 0.15 |
| Indole-3-carboxaldehyde | 146.0606 | 2.5 - 10000 | 0.9998 | 80 | 1.37 | 0.15 |
| Indole-3-carboxaldehyde-8C13 | 154.0874 | - | - | 80 | 1.37 | 0.15 |
| Indole-3-lactic acid | 206.0817 | 5 - 10000 | 0.9913 | 50 | 1.32 | 0.15 |
| Tryptamine | 161.1079 | 2.5 - 10000 | 0.9998 | 20 | 1.05 | 0.20 |
| 3-Hydroxyanthranilic acid | 154.0504 | 500 - 10000 | 0.9963 | 40 | 0.78 | 0.20 |
| Indole-3-acetamide | 175.0871 | 2.5 - 10000 | 0.9997 | 30 | 1.24 | 0.20 |

Note: CE: collision energy. RT: retention time. Linear dynamic range and R<sup>2</sup> were obtained using indole-3-carboxaldehyde-8C13 as the internal standard. Each concentration was measured in triplicate.

Major mass spectrometer parameters were as follows: spray voltage at 2750 V, sheath gas at 60 arb, aux gas at 10 arb, sweep gas at 1 arb, ion transfer tube temperature at 325 ℃, and vaporizer temperature at 400 ℃. The isolation window was set to 0.4 m/z without isolation offset. The collision energy was fixed; the normalized collision energy for each compound is listed in **Table 1**. PRM acquisition for each compound was scheduled, with optimized collision energy. The orbitrap resolution was set to 15,000, with the scan range mode set to auto. The AGC target was standard, the maximum injection time mode was auto, the intensity threshold was 1e4, dynamic exclusion was 6 seconds, and microscan was set to 1. Data was collected in profile mode, and quantitation was achieved using Skyline.^30^ The data is available at the Metabolomics Workbench.^31^ Study ID for *B. uniformis*: ST003344; doi: 10.21228/M8RJ9N. Study ID for ME/CFS fecal samples: ST003346; doi: 10.21228/M8RJ9N.

## RESULTS AND DISCUSSION

### Metabolomic Profiling of Bacteria with Different AhR-Activating Potencies

In a previous study, *S. epidermidis* and *E. faecium* were isolated from a human host and cultured. The culture supernatant of *S. epidermidis* exhibited high AhR-activating potency, whereas *E. faecium* showed low potency. Full extracts of their culture supernatants and HPLC fractions were analyzed using quadrupole-time-of-flight mass spectrometry in positive ion mode and open-source software.^29^

In this study, metabolomics profiling of these supernatants and fractions was performed on a quadrupole-orbitrap instrument in both positive and negative ion modes. Data were analyzed using Compound Discoverer with indole as a target structure during the database search to maximize the examination of indole derivatives in these two bacteria. **Figure 1** compares identifications and mapped mass variants between *S. epidermidis* and *E. faecium* samples: (A) full extracts and (B) SF12, SF13, and SF14 of the two bacteria culture supernatants (SF12, SF13, and SF14 were 30-second fractionations carrying a significant percentage of AhR-activating potency for the *S. epidermidis* supernatant; corresponding fractions of *E. faecium* were used as controls).^29^ Over 20 indole compounds in tryptophan pathways^20^ as well as indole-3-acetate conjugates were identified (**Figure 1**).

**Figure 1.**
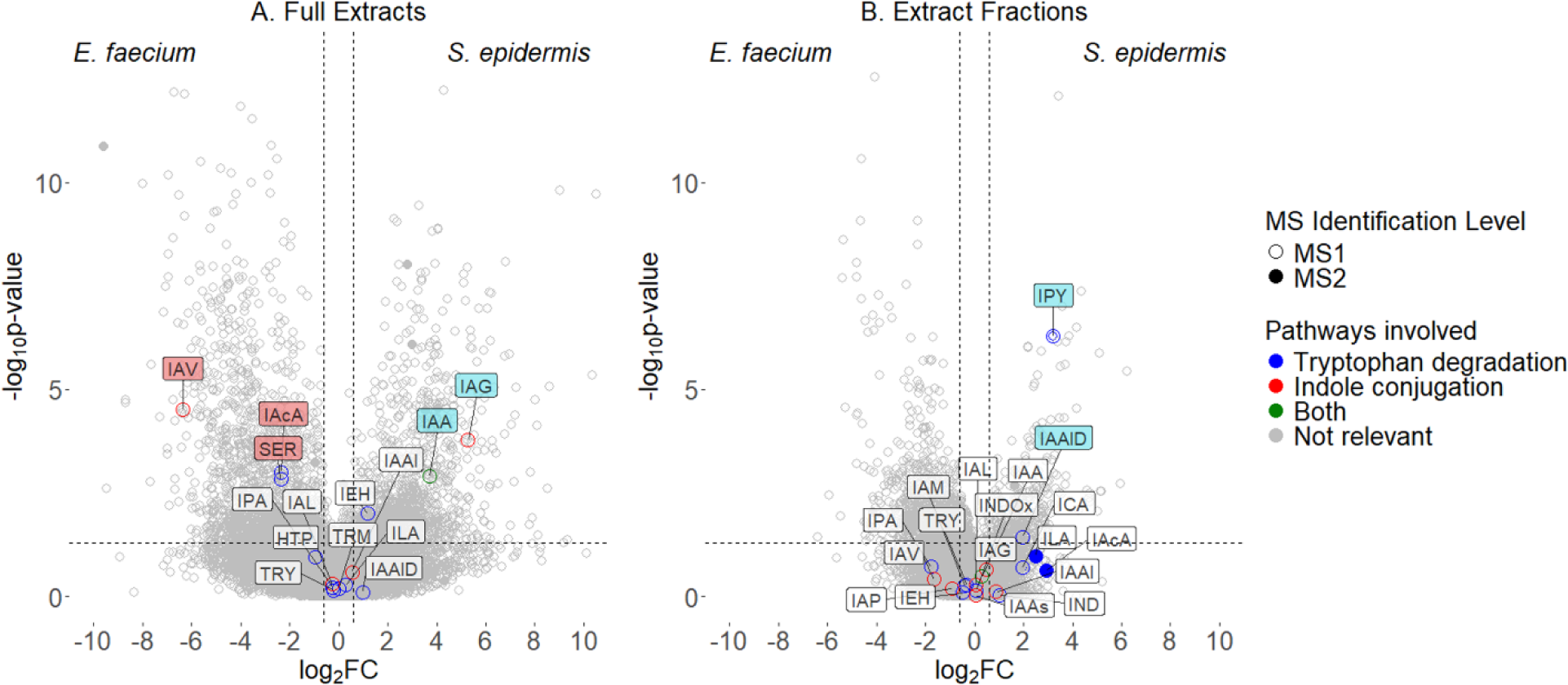
Relative amounts of metabolites in culture supernatants of *S. epidermidis* versus *E. faecium* strains. The *S. epidermidis* strain has a high potency for activating the aryl hydrocarbon receptor, whereas *E. faecium* has a low potency. Identification and label-free quantitation of metabolites were based on metabolomic profiling data using both positive and negative electrospray ionization. A: Volcano plot for full extracts; B: Volcano plot for extract fractions of high activating potential. P-value cutoff was set to 0.05, and fold change was set to 2. HTP: 5-Hydroxy-tryptophan; IAA: Indole-3-acetic acid; IAAlD: Indole-3-acetaldehyde; IAAl: Indole-3-acetyl alanine; IAAs: Indole-3-acetyl aspartic acid; IAcA: Indole-3-acrylic acid; IAG: Indole-3-acetyl glutamic acid; IAL: Indole-3-acetyl leucine; IAM: Indole-3-acetamide; IAP: Indole-3-acetyl phenylalanine; IAV: Indole-3-acetyl valine; ICA: Indole-3-aldehyde; IEH: Indole-3-ethanol; ILA: Indole-3-lactic acid; IPA: Indole-3-propionic acid; IPY: Indole-3-pyruvic acid; IND: Indole; INDOx: Indoxyl; SER: Serotonin; TRM: Tryptamine; TRY: Tryptophan.

Indole-3-acetic acid and indole-3-ethanol in the indole pathway had confident concentration increase in the full extracts of *S. epidermidis* over *E. faecium* (**Figure 1A**). Fractionation of culture supernatants observed additional indole derivatives like indole-3-pyruvic acid and indole-3-acetaldehyde, which had higher concentrations for *S. epidermidis* compared to the corresponding fractions of *E. faecium* (**Figure 1B**). A confident fold change for indole-3-acetaldehyde was not obtained from the metabolomic profiling of the full extracts. Additional metabolites from bacterial tryptophan degradation (**Scheme S1**) were identified but not confidently quantified by the DDA method. Many of these indole compounds and other tryptophan metabolites are known agonists of AhR.^17,32^

Multiple amino acid conjugates of indole-3-acetic acid in the superpathway of indole-3-acetate conjugate biosynthesis (**Scheme S2**) were also identified. Interestingly, some showed significant differences in concentration when comparing *S. epidermidis* and *E. faecium*. The supernatant of *S. epidermidis* had a higher level of glutamic acid conjugate, whereas *E. faecium* had a higher concentration of valine conjugate (**Figure 1A**). Indole-3-acetic acid is an important plant hormone. Its amino acid conjugates, which have reduced biological activity, serve as storage forms of indole-3-acetic acid, helping the regulation of a controlled supply of the active hormone.^33^ However, little is known about microbial amino acid conjugation of indole-3-acetic acid and whether these conjugates play a role in regulating the host AhR pathway.

Overall, although many indole compounds in the kynurenine, serotonin, and indole pathways were identified, their quantitation was below the level of statistical confidence (**Figure 1**). Thus, methods with improved figures of merit are needed to better examine microbial interaction of indole derivatives with the human host via the AhR pathway.

### Targeted Metabolomics for Comparing Metabolic Diversity of Bacterial Tryptophan Metabolites among Genera, Species, and Strains

To address the profiling method limitations, we employed a targeted metabolomics approach. Targeted quantitation allows for the specific measurement of metabolites of interest, providing improved sensitivity, accuracy, and precision. To accommodate the wide concentration range of select metabolites, varying by three orders of magnitude, full supernatant extracts were reconstituted and analyzed as 5-fold diluted or 5-fold concentrated solutions relative to the original supernatants, aligning with the established calibration curves. Notably, certain target indole compounds—including indole-3-acetonitrile, indole-3-butyrate, 5,11-dihydroindolo[3,2-b]carbazole, and methyl 2-(1H-indole-3-carbonyl)thiazole-4-carboxylate—remained undetectable even in 5X concentrated supernatant samples.

Quantified tryptophan catabolites in bacterial culture supernatant extracts—tryptamine, indole-3-acrylate, indole-3-ethanol, indole-3-propionate, indole-3-acetate, indole-3-aldehyde, indole-3-lactate, N-formylkynurenine, and kynurenine—reveal metabolic diversity in tryptophan catabolism during cell culture. Relative concentrations, expressed as percentages, accounting for variations in bacterial culturing process like uneven cell density and activity during overnight growth, provides a consistent basis for analysis.

It is essential to note that metabolite quantitation using calibration curves only produces relative amounts, especially when metabolites differ in structure (and likely in ionization efficiency) from internal quantitation standards. Therefore, percentage concentrations are preferred for depicting metabolic differences in pathway among different bacteria, which enables flexibility in the selection of quantitation standards.

We constructed two distribution plots, with and without N-formylkynurenine and kynurenine (**Figure 2A**, **Table S3** and **Figure 2B, Table S4**), to separate indole derivatives associated with the bacterial indole pathway from these early kynurenine pathway metabolites. N-formylkynurenine and kynurenine, shared by bacteria and humans, dominate many bacterial culture supernatants (**Figure 2A**). By excluding these metabolites, significant variations in tryptophan catabolite diversity within the bacterial indole pathway became evident (**Figure 2B**), underscoring the value of nuanced analysis in analyzing metabolic pathways.

**Figure 2.**
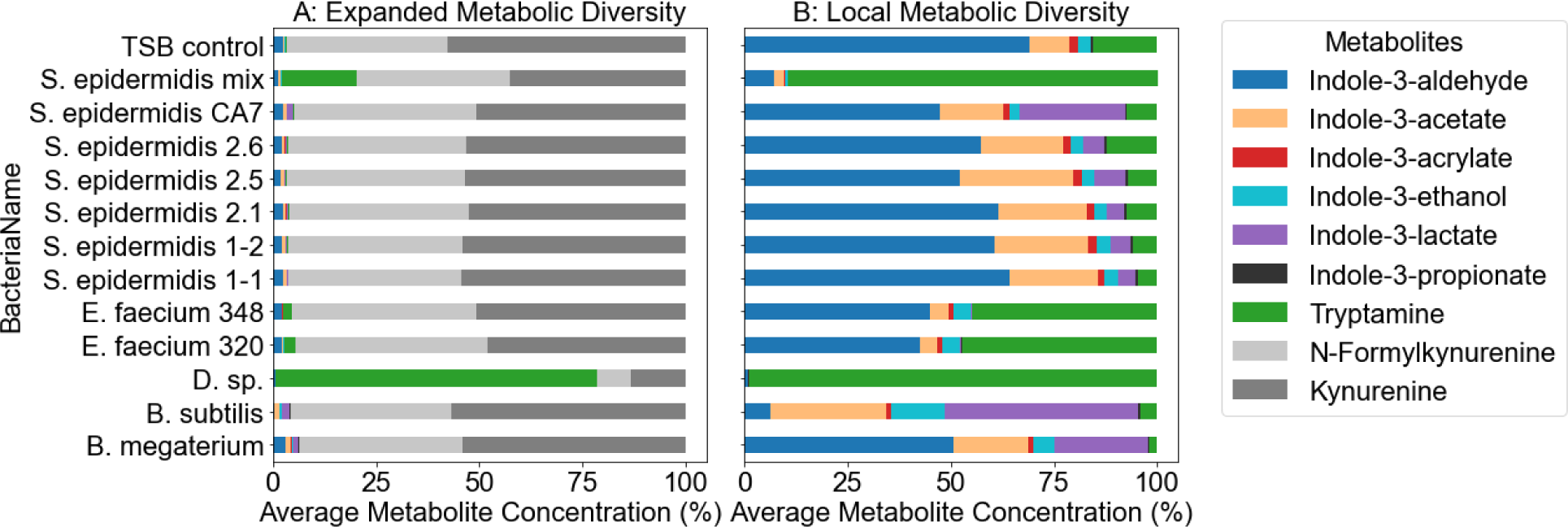
Relative Concentrations of Indole Derivatives for Characterizing Metabolic Diversity in the Bacterial Indole Pathway. A: Distributions of relative concentrations of tryptophan catabolites in bacteria of different genera, species, and strains. Catabolites included compounds from bacterial indole pathway and kynurenine pathway shared by human and bacteria. B: Distributions of indole derivatives as measures for local metabolic diversity in the indole pathway of different bacteria.

The principal component analysis (PCA) of these two sets of tryptophan catabolite relative concentrations yielded similar outcomes. However, compared to a broader analysis of tryptophan catabolite diversity (**Figure 3A**), examining the more specific indole derivatives (**Figure 3B**) revealed higher explained variances in Principal Component 1 (PC1) and Principal Component 2 (PC2). The broader analysis presented a PC1 of 53.1% and a PC2 of 23.6%, while the focused analysis on indole diversity showed a PC1 of 59.9% and a PC2 of 27.5%.

**Figure 3.**
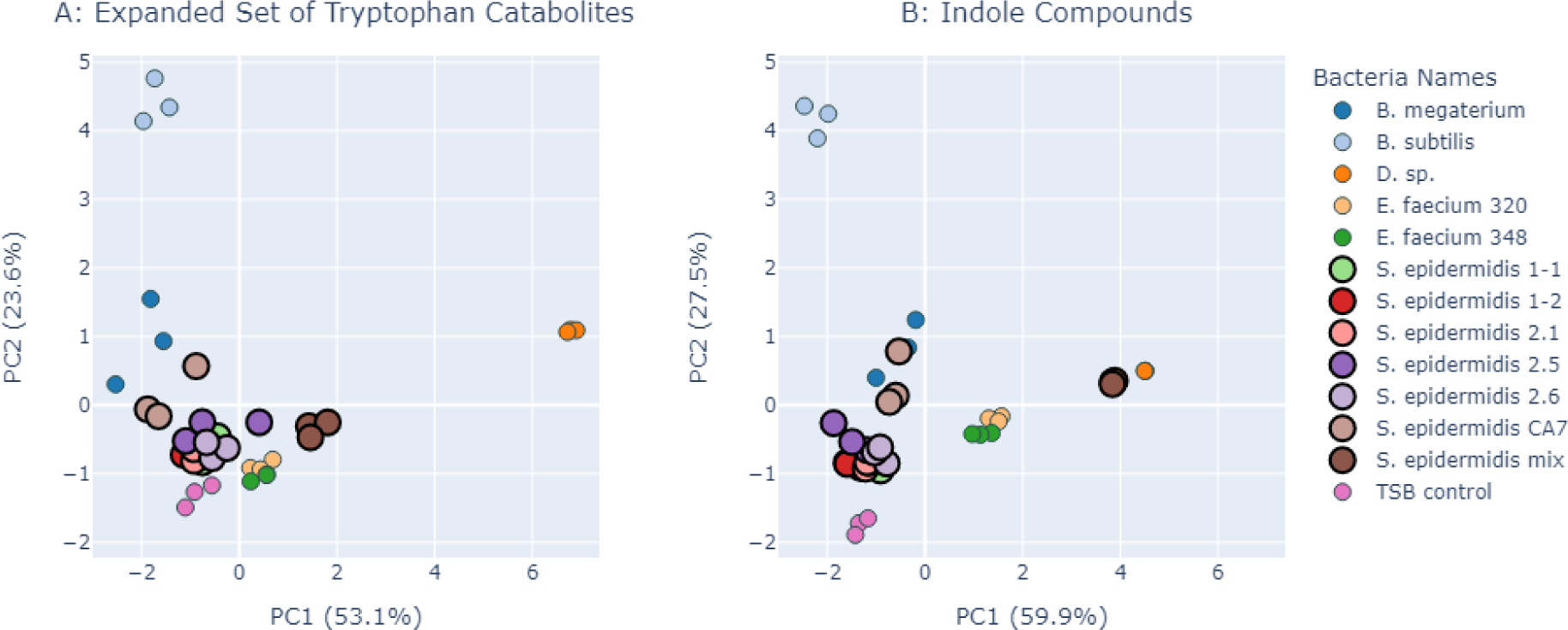
Metabolic Diversity of Tryptophan Metabolites for Bacteria Characterization. A: PCA of relative concentrations of tryptophan catabolites. B: PCA of relative concentrations of bacterial indole derivatives. Highlighted dots (in larger size) for *S. epidermis* bacteria showed potential utilization of relative concentrations of bacterial indole derivatives—local metabolic diversity—for the differentiation of different bacteria strains.

The use of relative concentrations for indole pathway metabolites facilitated a more cohesive clustering of bacteria across various genera, species, and strains (**Figure 3B**). In the PCA plot, the TSB control was distinctly separated from all bacterial samples, indicating clear differentiation. The two *E. faecium* strains formed a tight cluster, suggesting similar metabolic profiles. Different *S. epidermidis* strains, while positioned closely, showed slight separations, with a mix of sub-strains diverging from individual sub-strain contributors. This separation highlights potential metabolic interplay among sub-strains during culture.^5^ These findings prompt further research into the analytical utility of local metabolic diversity in indole derivatives for differetiating bacterial strains and classifying human microbiota samples for disease research. Given the large sample sizes preferred for these analyses, there is a need to improve sample throughput of targeted metabolomics techniques for practicality.

### Development of a 4-Minute Method of Fast Targeted Metabolomics to Improve Sample Throughput

To develop a fast, highly selective targeted metabolomics method, we opted for LC-PRM MS over LC-SRM MS. The low resolution in ion selection with SRM poses limitations for analyzing metabolomics samples,^27^ an issue that becomes more pronounced with shortened LC run times in high-throughput analyses. PRM addresses this problem effectively by complementing reduced chromatographic separation with high-resolution analysis of fragment ions in the gas phase. Additionally, PRM does not require pre-selection and optimization of specific fragment ions. Instead, it utilizes post-acquisition data mining and allows retrospective analysis to detect newly identified metabolites or unforeseen contaminants in past data. During the fast LC-PRM method development, we focused on internal standard selection and method reproducibility.

Requirements for isotopic quantitation references in metabolomics differ significantly from those in proteomics. In proteomics, the pairwise use of isotopically labeled peptides for quantitation is practical, necessitating the co-elution of light and heavy peptides. This process requires more expansive isotopic labeling with ^13^C, ^15^N, and ^18^O to limit matrix effect variances. In contrast, the limited availability and additional costs of isotopic metabolites for pairwise use in targeted metabolomics render this approach less practical. Moreover, characterizing the metabolic diversity of indole derivatives only requires relative quantitation of the compounds, making the pairwise use of isotopic standards unnecessary.

Normalization-based quantitation, utilizing a few isotopic standards, including cost-effective deuterium-based standards, was selected for the targeted quantitation of indole compounds. The accuracy and precision of this approach hinge on the reproducibility of elution times for both quantitation standards and target analytes, rather than their coelution. Ensuring consistent run-to-run elution times is increasingly critical as reduced separation times are employed to enhance sample throughput. During the assessment of method reproducibility and robustness, through the injection of varying volumes of a standard mixture, low coefficients of variation were observed for both quantification (CV% = 5.4%) and retention time (CV% = 0.3%, **Table S5**).

The quantitation dynamic ranges of indole compounds based on the internal standards NFK-C13, IPA-2D, and ICA-8C13 were evaluated for isotopic interference (**Figure S2**). NFK-C13, used in the 28-minute PRM method, exhibited a limited dynamic range due to significant isotopic overlap; the natural C-13 peak represented about 11.75% of the C-12 peak of non-labeled N-formylkynurenine, leading to variable effects on the NFK-C13 monoisotopic intensity across samples (**Figure S3**). In comparison, IPA-2D demonstrated an improved dynamic range (R^2^ = 0.9999) with a 2 Da separation between non-labeled and isotopic forms, reducing isotopic overlap to 0.56%. Similarly, ICA-8C13, with an 8 Da mass separation, also showed a satisfactory dynamic range (R^2^ = 0.9966, **Figure S3**). Consequently, ICA-8C13 and IPA-2D were selected as isotopic internal standards for quantifying indole compounds using the fast PRM method. The use of an OT-2 robot for preparing calibration solutions and spiking quantitation references was integral to the fast method for building calibration curves. Twelve concentration levels of each reference compound, with eight replicates for target metabolites, were utilized to build calibration curves, and 0.50 µM of the standard was spiked in. Calibration curves for 13 target compounds exhibited strong linearity (R² > 0.99, **Figure S4**).

Sample throughput is crucial for exploiting the metabolic diversity of indole derivatives in the characterization and analysis of bacterial isolates from human hosts and patient fecal samples. Although several methods for the targeted quantitation of tryptophan metabolites have been reported, they usually require about 7-15 minutes per sample.^21–27^ Consequently, analyzing a 96-well plate in triplicate typically spans two days, leading to prolonged instrument run times, sample stability concerns, and substantial solvent consumption. The fast targeted metabolomics method had a rate of 4 minutes per sample (2.5-minute gradient) and achieved a 2 to 4-fold improvement in the sample throughput. Compared with the 28-minute method, the fast method has satisfactory MS resolution (15k for the 28-minute method and 20k for the 4-minute method at 200 *m/z*). A similar peak separation was achieved by the fast method based on high-resolution extracted ion chromatogram (**Figure S5**).

### Metabolic Diversity of Indole Derivatives in Bacteroides uniformis Strains

*B. uniformis*, a clinically significant species within the Bacteroides genus, are anaerobic, fiber-degrading bacteria that colonize the gut during early infancy, contributing to gut barrier integrity through butyrate and γ-aminobutyric acid production.^6–8,34^ *B. uniformis* strains from 12 ME/CFS patients and 12 healthy controls were cultured, and their culture supernatants were desalted and concentrated 20-fold for analysis. Indole derivatives were measured using the developed fast targeted metabolomics method (**Table S6**). PCA on the relative concentrations of these metabolites identified two major components (PC1, 68.9%; PC2, 18.7%) that accounted for 87.6% of the explained variance, distinguishing two distinct *B. uniformis* clusters (**Figure 4A**).

**Figure 4.**
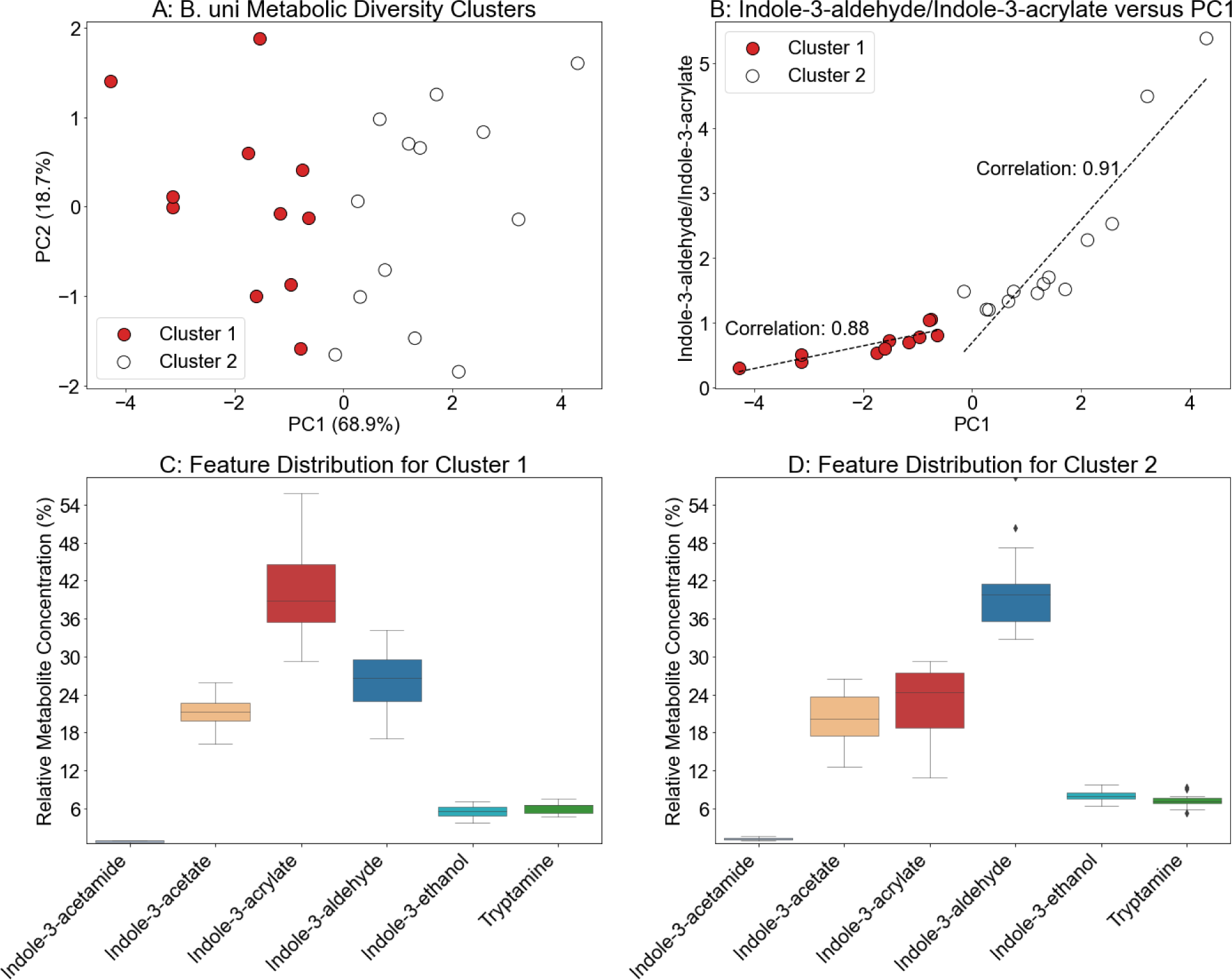
Metabolic Diversity of Indole Derivatives for Strain Differentiation of *Bacteroides uniformis*. A: PCA identified two clusters (Cluster 1 in solid red dots, Cluster 2 in open dots) of B. uni bacteria strains. PC1 and PC2 represented over 86% of the explained variation in the data. Twenty-four *B. uni* strains were cultured in biological triplicate on the same well plate. Each bacterial culture supernatant was measured in triplicate to obtain metabolite concentrations. B: the ratio of relative concentrations for indole-3-aldehyde to indole-3-acrylate had strong correlation with PC1, with correlation coefficients for Cluster 1 = 0.88 and Cluster 2 = 0.91, respectively. C and D: Feature distributions for Clusters 1 and 2, identifying flipping ratios for indole-3-aldehyde and indole-3-acrylate in two clusters.

Notable metabolic diversity was evident between the two clusters, particularly in the contributions of indole-3-acrylate and indole-3-aldehyde (**Figures 4B, 4C, 4D**). The relative concentration ratios of these compounds strongly correlated with PC1, with correlation coefficients of 0.88 and 0.91 for Clusters 1 and 2, respectively, underscoring their distinct roles in the indole pathway. The ability of genetically identical bacteria to manifest variations in metabolic phenotype is likely attributed to interactions with the complex gut environment.^4^ It has been reported that gut microbiota composition and activity influence the host metabolic phenotype and may relate to diseases.^2^ Given that indole-3-aldehyde and indole-3-acrylate are more reactive compared to other tryptophan catabolites in the indole pathway, investigating their potential influence on the metabolic adaptation of *B. uniformis* strains within the human gut microbiota presents a compelling area of research. It should be noted, however, that a direct correlation between *B. uniformis* strains and ME/CFS was not obtained.

### Machine Learning to Classify Metabolic Diversity of Indole Derivatives in the ME/CFS Gut Microbiome

Exploring the link between microbial metabolites in human samples and health conditions requires extensive studies with large patient cohorts due to the inherent complexity and heterogeneity of the samples. To achieve statistically significant results, it is essential to conduct replicated analyses on a large number of samples. The fast targeted metabolomics method developed was used to analyze fecal samples from 10 ME/CFS patients and 10 healthy controls, each in triplicate. Samples were lyophilized, and an equal mass was extracted from each, subsequently desalted, and reconstituted for analysis. Among the target metabolites listed in **Table 1**, five compounds in the bacterial indole pathway were quantified under the experimental conditions: indole-3-ethanol, indole-3-propionate, indole-3-acetate, indole-3-aldehyde, and tryptamine (**Table S7**).

One significant advantage of using metabolite concentration distribution—relative concentrations—to characterize fecal samples is that initial sampling can vary significantly due to high sample heterogeneity. Therefore, common measures of metabolites based on fecal weight are questionable. Additionally, many metabolite targets lack isotopic quantitation standards, making their absolute quantitation challanging.

PCA of the concentration distributions of the five quantified indole compounds did not distinguish ME/CFS samples from control samples into separate clusters. Instead, it organized all samples into three mixed clusters (**Figure 5A**), encompassing both patient and control samples. This outcome, along with the varying relative concentrations of indole derivatives (**Figure 5B**), highlights the complex interplay in the production and metabolism of indole compounds within the human gut microbiome and their interactions with the host. It suggests that the metabolic networks involved are more complex than straightforward disease correlations. Thus, advanced data analysis techniques are required to uncover the disease insights offered by the fast targeted metabolomics of the metabolic diversity of indole derivatives.

**Figure 5.**
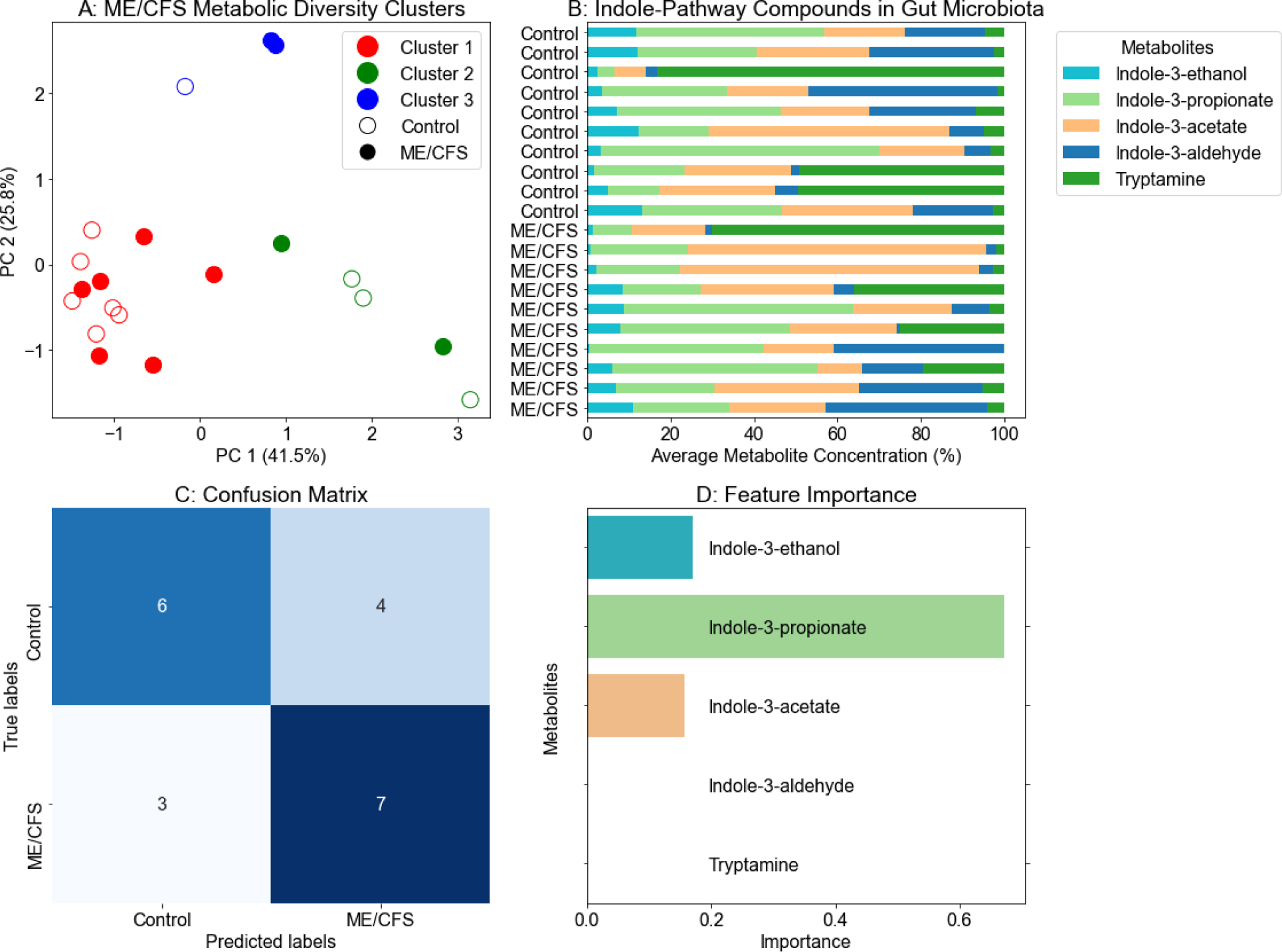
Linking metabolic diversity of bacterial indole derivatives with ME/CFS. A: PCA of relative concentrations of indole derivatives in ME/CFS (open circles, n=10) and controls (solid circles, n=10) fecal samples. Samples are grouped into three clusters (red, green, and blue). Each fecal sample was extracted in triplicate, and fast LC-PRM MS measured each replicate in triplicate. Average concentrations were used to calculate the relative concentration (%) for each sample. B: Percentage distributions of indole derivatives in fecal samples. C: Confusion matrix for sample classification. D: Relative importance of features for the AdaBoost-LOOCV model.

Limited by the cohort size, AdaBoost was used to develop a model with 10 estimators, a learning rate of 0.5, and a decision tree depth of 2 for classifying fecal samples. This model underwent validation through leave-one-out cross-validation (LOOCV), achieving a mean LOOCV accuracy of 0.65 (Control: precision of 0.67, recall of 0.60, F1-score of 0.63; ME/CFS: precision of 0.64, recall of 0.70, F1-score of 0.67), indicating moderate classification success for distinguishing between ME/CFS and control samples (**Figure 5C**). Indole-3-propionate emerged as the most important feature in model training, with indole-3-ethanol and indole-3-acetate also identified as significant features (**Figure 5D**). Notably, indole-3-ethanol exhibited low relative concentrations across all samples. Despite the minimal feature importance of indole-3-aldehyde and tryptamine, their inclusion was crucial for the model to observe classification success. While indole-3-aldehyde was a determining feature in differentiating *B. uniformis* strains, its relatively high chemical reactivity could result in its instability in complex fecal samples. In contrast, the relative low reactivity of indole-3-propionate, indole-3-ethanol, and indole-3-acetate could contribute to their robustness to merge as features in machine learning. Improving the AdaBoost-LOOCV model for ME/CFS diagnostic and therapeutic applications can benefit from a larger cohort size and the inclusion of additional local metabolic diversity beyond the bacterial indole pathway.

## CONCLUSION

This work proposes to use local metabolic diversity for characterizing bacterial strains and classifying microbiome samples. A fast, high-resolution LC-PRM MS method with a throughput rate of 4 minutes per sample was developed, significantly enhancing sample throughput for targeted metabolomics of indole derivatives. The increased sample throughput facilitates large-scale analysis of complex and heterogeneous samples. Targeted metabolomic analysis of culture supernatants from 24 strains of *B. uniformis* revealed a characteristic metabolic difference in indole pathway derivatives. The ratio of indole-3-acrylate to indole-3-aldehyde was reversed between two distinct *B. uniformis* groups. Furthermore, indole compounds in ME/CFS and control fecal samples were measured, and results were analyzed using AdaBoost for machine learning and LOOCV for validation, achieving moderate success in distinguishing ME/CFS from control samples. Key features for the classification included indole-3-propionate, indole-3-ethanol, and indole-3-acetate. Therefore, the local metabolic diversity in the bacteria indole pathway is a promising biomarker candidate for differentiating bacterial strains and classifying ME/CFS samples. Enhancements to the AdaBoost-LOOCV model can use larger cohort sizes and include additional pathways to further elevate the diagnostic and therapeutic utilities of the local metabolic diversity in studying microbiome contributions to ME/CFS and other immune-related conditions.

## Acknowledgments

This work was supported by the National Institutes of Health (1U54NS105539-01).

## Data Availability Statement

This study is available at the NIH Common Fund’s National Metabolomics Data Repository (NMDR) website, the Metabolomics Workbench, https://www.metabolomicsworkbench.org, supported by NIH U2C-DK119886 and OT2-OD030544 grants. Study ID for 28-minute method quantitation of indole compounds: ST002308, doi: 10.21228/M8G13Q. Study ID for 4-minute method quantitation of indole compounds in *B. uniformis*: ST003344, doi: 10.21228/M8RJ9N. Study ID for fast targeted metabolomics of ME/CFS fecal samples: ST003346, doi: 10.21228/M8RJ9N. The data can be accessed directly via corresponding Project DOI.

## Supporting Information

Table S1, twelve bacteria strains used for the 28-min method; Table S2, selected indole compounds for targeted quantitation using scheduled LC-PRM; Table S3, tryptophan metabolites in bacterial cell culture supernatant; Table S4, quantified indole compounds in different bacteria; Table S5, reproducibility test of standard with five replicates for the 4-min method; Table S6, quantified indole derivatives of 24 *Bacteroides uniformis* strains; Table S7, quantified compounds of the bacterial indole pathway in fecal samples; Figure S1. calibration curves for the 28-min method; Figure S2, structure and isotope overlap of light and heavy tryptophan catabolites; Figure S3, calibration curves for kynurenine using different internal standards; Figure S4, calibration curves for the 4-min method; Figure S5, screenshots of ion chromatogram and skyline ion chromatogram of the old 28-min method and the new 4-min method; Scheme S1, Bacterial tryptophan degradation; Scheme S2, Superpathway of indole-3-acetate conjugate biosynthesis.

