## Supplemental Information for "Fast Targeted Metabolomics for Analyzing Metabolic Diversity of Bacterial Indole Derivatives in ME/CFS Gut Microbiome"

**Table S1.** Twelve bacteria strains used for the 28-min method.

| Species | Location | Gram+/- | Media | Aerobic/Anaerobic |
|---|---|---|---|---|
| *Enterococcus faecium* 348 | patient gut | + | TSB | Aerobic |
| *Dermabacter* sp. | ATCC | + | TSB | Aerobic |
| *Bacillus subtilis* | healthy skin | + | TSB | Aerobic |
| *Bacillus megaterium* | healthy oral | + | TSB | Aerobic |
| *Staphylococcus epidermidis* mix | healthy gut | + | TSB | Aerobic |
| *Enterococcus faecium* 320 | healthy gut | + | TSB | Aerobic |
| *Staphylococcus epidermidis* CA7 | healthy skin | + | TSB | Aerobic |
| *Staphylococcus epidermidis* 1-1 | sub-strain of Staphylococcus epidermidis mix | + | TSB | Aerobic |
| *Staphylococcus epidermidis* 1-2 | sub-strain of Staphylococcus epidermidis mix | + | TSB | Aerobic |
| *Staphylococcus epidermidis* 2-1 | sub-strain of Staphylococcus epidermidis mix | + | TSB | Aerobic |
| *Staphylococcus epidermidis* 2-5 | sub-strain of Staphylococcus epidermidis mix | + | TSB | Aerobic |
| *Staphylococcus epidermidis* 2-6 | sub-strain of Staphylococcus epidermidis mix | + | TSB | Aerobic |

**Table S2.** Selected indole compounds for targeted quantitation using scheduled LC-PRM.

| Compound Name | Theoretical *m/z* | Linear Range (nM) | $R^2$ | Normalized CE (%) | RT (min) | RT Window (min) |
|---|---|---|---|---|---|---|
| N-formylkynurenine-C13 | 238.0909 | - | - | 30 | 4.64 | 0.40 |
| N-formylkynurenine | 237.0870 | 0.5 - 1000 | 0.9982 | 30 | 4.64 | 0.40 |
| Kynurenine | 209.0926 | 0.5 - 1000 | 0.9969 | 30 | 4.34 | 0.40 |
| Indole-3-acetonitrile | 157.0766 | 100 - 2000 | 0.9936 | 80 | 9.76 | 0.40 |
| Indole-3-acrylic acid | 188.0712 | 0.5 - 1000 | 0.9987 | 50 | 8.95 | 0.40 |
| Indole-3-ethanol | 162.0919 | 0.5 - 1000 | 0.9986 | 40 | 8.39 | 0.40 |
| Indole-3-propionic acid | 190.0868 | 0.5 – 1000 | 0.9985 | 60 | 9.23 | 0.40 |
| Indole-3-acetic acid | 176.0712 | 0.5 - 1000 | 0.9979 | 90 | 8.35 | 0.40 |
| Indole-3-butyric acid | 204.1025 | 0.5 - 1000 | 0.9971 | 80 | 9.89 | 0.40 |
| Indole-3-carboxaldehyde | 146.0606 | 0.5 - 1000 | 0.9979 | 80 | 8.06 | 0.40 |
| Indole-3-lactic acid | 206.0817 | 0.5 - 1000 | 0.9985 | 50 | 7.64 | 0.40 |
| Tryptamine | 161.1079 | 1 - 1000 | 0.9939 | 20 | 5.83 | 0.40 |
| Indole | 118.0657 | 0.5 -2000 | 0.9945 | 110 | 10.23 | 0.40 |
| 5,11-dihydroindolo[3,2-b]carbazole | 257.1079 | 5 - 1000 | 0.9962 | 90 | 13.12 | 0.40 |
| methyl 2-(1H-indole-3-carbonyl) thiazole-4-arboxylate | 287.0490 | 50 - 2000 | 0.9921 | 90 | 11.54 | 0.40 |

Note: CE: collision energy. RT: retention time. Linear dynamic range and $R^2$ were obtained using N-formylkynurenine-C13 as the internal standard.

**Table S3.** Tryptophan metabolites in bacterial cell culture supernatant.

| IaId | IAA | IA | IE | ILA | IPA | TRYA | FKYN | KYN | Bacteria Name |
|---|---|---|---|---|---|---|---|---|---|
| 3.16 | 1.33 | 0.07 | 0.3 | 2.02 | 0.01 | 0.15 | 39.93 | 53.02 | B. megaterium |
| 3.19 | 1.06 | 0.07 | 0.32 | 1.36 | 0.02 | 0.24 | 37.82 | 55.92 | B. megaterium |
| 3.36 | 1.06 | 0.11 | 0.34 | 1.05 | 0.02 | 0.01 | 40.45 | 53.61 | B. megaterium |
| 0.23 | 1.23 | 0.07 | 0.48 | 2.19 | 0.03 | 0.19 | 39.13 | 56.45 | B. subtilis |
| 0.28 | 1.15 | 0.05 | 0.56 | 1.87 | 0.03 | 0.07 | 38.51 | 57.48 | B. subtilis |
| 0.3 | 1.26 | 0.06 | 0.62 | 2.04 | 0.02 | 0.3 | 38.92 | 56.47 | B. subtilis |
| 0.55 | 0.09 | 0.03 | 0.08 | 0 | 0.01 | 76.83 | 8.31 | 14.11 | D. sp. |
| 0.6 | 0.1 | 0.03 | 0.09 | 0 | 0.01 | 77.87 | 7.99 | 13.32 | D. sp. |
| 0.68 | 0.11 | 0.03 | 0.1 | 0 | 0.01 | 78.63 | 8.03 | 12.42 | D. sp. |
| 1.99 | 0.22 | 0.06 | 0.21 | 0.02 | 0.01 | 2.17 | 44.39 | 50.92 | E. faecium 320 |
| 2.5 | 0.22 | 0.05 | 0.24 | 0 | 0.01 | 2.68 | 47.4 | 46.89 | E. faecium 320 |
| 2.51 | 0.27 | 0.06 | 0.25 | 0.02 | 0.01 | 2.97 | 47.3 | 46.61 | E. faecium 320 |
| 1.89 | 0.19 | 0.07 | 0.19 | 0.01 | 0.01 | 1.79 | 45.27 | 50.59 | E. faecium 348 |
| 2.1 | 0.2 | 0.06 | 0.18 | 0 | 0.01 | 2.32 | 42.44 | 52.69 | E. faecium 348 |
| 2.33 | 0.21 | 0.07 | 0.21 | 0.01 | 0.01 | 2.17 | 45.7 | 49.29 | E. faecium 348 |
| 2.8 | 0.78 | 0.06 | 0.13 | 0.14 | 0.02 | 0.27 | 42.76 | 53.04 | S. epidermidis 1-1 |
| 2.28 | 0.83 | 0.06 | 0.13 | 0.16 | 0.02 | 0.13 | 40.01 | 56.38 | S. epidermidis 1-1 |
| 2.34 | 0.82 | 0.06 | 0.14 | 0.16 | 0.02 | 0.16 | 42.89 | 53.41 | S. epidermidis 1-1 |
| 2.05 | 0.82 | 0.08 | 0.12 | 0.17 | 0.02 | 0.24 | 41.31 | 55.19 | S. epidermidis 1-2 |
| 2.04 | 0.78 | 0.07 | 0.12 | 0.17 | 0.02 | 0.16 | 43.23 | 53.41 | S. epidermidis 1-2 |
| 2.53 | 0.86 | 0.08 | 0.13 | 0.17 | 0.02 | 0.26 | 42.74 | 53.21 | S. epidermidis 1-2 |
| 2.4 | 0.81 | 0.07 | 0.12 | 0.16 | 0.02 | 0.32 | 44.8 | 51.3 | S. epidermidis 2.1 |
| 2.25 | 0.78 | 0.07 | 0.11 | 0.17 | 0.02 | 0.28 | 42.77 | 53.55 | S. epidermidis 2.1 |
| 2.53 | 0.88 | 0.07 | 0.13 | 0.17 | 0.02 | 0.26 | 43.13 | 52.81 | S. epidermidis 2.1 |
| 1.04 | 0.72 | 0.06 | 0.08 | 0.25 | 0.01 | 0.18 | 42.48 | 55.18 | S. epidermidis 2.5 |
| 2.24 | 0.99 | 0.08 | 0.12 | 0.23 | 0.02 | 0.41 | 45.24 | 50.67 | S. epidermidis 2.5 |
| 2.12 | 1 | 0.06 | 0.13 | 0.22 | 0.02 | 0.14 | 41.76 | 54.54 | S. epidermidis 2.5 |
| 2.23 | 0.82 | 0.07 | 0.12 | 0.25 | 0.02 | 0.46 | 43.31 | 52.72 | S. epidermidis 2.6 |
| 2.48 | 0.73 | 0.06 | 0.13 | 0.14 | 0.02 | 0.46 | 43.09 | 52.89 | S. epidermidis 2.6 |
| 1.81 | 0.7 | 0.07 | 0.11 | 0.18 | 0.02 | 0.47 | 43.16 | 53.49 | S. epidermidis 2.6 |
| 1.47 | 0.56 | 0.06 | 0.1 | 0.04 | 0.01 | 18.65 | 38.13 | 40.98 | S. epidermidis mix |
| 1.22 | 0.5 | 0.05 | 0.09 | 0.04 | 0.01 | 16.98 | 36.17 | 44.93 | S. epidermidis mix |
| 1.65 | 0.49 | 0.06 | 0.1 | 0.03 | 0.02 | 18.47 | 37.58 | 41.61 | S. epidermidis mix |
| 1.98 | 0.75 | 0.07 | 0.11 | 1.51 | 0.02 | 0.35 | 42.14 | 53.06 | S. epidermidis CA7 |
| 2.82 | 0.88 | 0.08 | 0.14 | 1.24 | 0.02 | 0.46 | 46.85 | 47.51 | S. epidermidis CA7 |
| 2.63 | 0.8 | 0.08 | 0.1 | 1.22 | 0.02 | 0.33 | 43.4 | 51.41 | S. epidermidis CA7 |
| 2.35 | 0.34 | 0.07 | 0.1 | 0 | 0.03 | 0.56 | 38.08 | 58.48 | TSB control |
| 2.22 | 0.36 | 0.08 | 0.09 | 0 | 0.03 | 0.67 | 39.11 | 57.43 | TSB control |
| 2.77 | 0.33 | 0.08 | 0.11 | 0 | 0.03 | 0.39 | 38.85 | 57.45 | TSB control |

Note: IA: Indole-3-acrylic acid; IAA: Indole-3-acetic acid; Iald: Indole-3-aldehyde; IE: Indole-3-ethanol; ILA: Indole-3-lactic acid; IPA: Indole-3-propionic acid; KYN: Kynurenine; FKYN: N-Formylkynurenine; TRYA: Tryptamine.

**Table S4.** Quantified indole compounds in different bacteria.

| Iald | IAA | IA | IE | ILA | IPA | TRYA | Bacteria Name |
|---|---|---|---|---|---|---|---|
| 44.92 | 18.82 | 1.04 | 4.22 | 28.68 | 0.16 | 2.16 | B. megaterium |
| 50.94 | 16.94 | 1.13 | 5.07 | 21.78 | 0.24 | 3.90 | B. megaterium |
| 56.58 | 17.81 | 1.76 | 5.77 | 17.64 | 0.27 | 0.17 | B. megaterium |
| 5.18 | 27.92 | 1.65 | 10.88 | 49.56 | 0.58 | 4.23 | B. subtilis |
| 6.98 | 28.69 | 1.33 | 14.05 | 46.66 | 0.65 | 1.63 | B. subtilis |
| 6.59 | 27.37 | 1.27 | 13.54 | 44.26 | 0.54 | 6.42 | B. subtilis |
| 0.70 | 0.12 | 0.03 | 0.10 | 0.00 | 0.01 | 99.03 | D. sp. |
| 0.77 | 0.12 | 0.03 | 0.11 | 0.00 | 0.01 | 98.95 | D. sp. |
| 0.85 | 0.14 | 0.04 | 0.12 | 0.00 | 0.01 | 98.84 | D. sp. |
| 42.51 | 4.76 | 1.21 | 4.57 | 0.46 | 0.23 | 46.27 | E. faecium 320 |
| 43.84 | 3.86 | 0.93 | 4.16 | 0.07 | 0.26 | 46.88 | E. faecium 320 |
| 41.19 | 4.39 | 1.01 | 4.16 | 0.34 | 0.22 | 48.68 | E. faecium 320 |
| 43.05 | 4.11 | 1.25 | 3.72 | 0.08 | 0.25 | 47.55 | E. faecium 348 |
| 46.45 | 4.26 | 1.34 | 4.19 | 0.27 | 0.25 | 43.25 | E. faecium 348 |
| 45.58 | 4.55 | 1.58 | 4.56 | 0.33 | 0.24 | 43.16 | E. faecium 348 |
| 66.71 | 18.50 | 1.45 | 3.09 | 3.28 | 0.53 | 6.41 | S. epidermidis 1-1 |
| 63.18 | 22.92 | 1.60 | 3.54 | 4.55 | 0.55 | 3.66 | S. epidermidis 1-1 |
| 63.34 | 22.19 | 1.74 | 3.69 | 4.24 | 0.57 | 4.23 | S. epidermidis 1-1 |
| 58.69 | 23.43 | 2.20 | 3.40 | 4.90 | 0.54 | 6.84 | S. epidermidis 1-2 |
| 60.81 | 23.33 | 2.06 | 3.52 | 5.05 | 0.56 | 4.67 | S. epidermidis 1-2 |
| 62.47 | 21.28 | 1.86 | 3.24 | 4.18 | 0.55 | 6.42 | S. epidermidis 1-2 |
| 61.44 | 20.73 | 1.91 | 3.08 | 4.21 | 0.52 | 8.11 | S. epidermidis 2.1 |
| 61.10 | 21.23 | 1.81 | 3.10 | 4.64 | 0.50 | 7.61 | S. epidermidis 2.1 |
| 62.39 | 21.65 | 1.65 | 3.20 | 4.12 | 0.55 | 6.43 | S. epidermidis 2.1 |
| 44.35 | 31.01 | 2.62 | 3.28 | 10.74 | 0.42 | 7.57 | S. epidermidis 2.5 |
| 54.88 | 24.11 | 1.85 | 2.88 | 5.72 | 0.50 | 10.10 | S. epidermidis 2.5 |
| 57.43 | 26.97 | 1.72 | 3.45 | 6.03 | 0.54 | 3.85 | S. epidermidis 2.5 |
| 56.31 | 20.63 | 1.69 | 3.07 | 6.34 | 0.49 | 11.50 | S. epidermidis 2.6 |
| 61.73 | 18.08 | 1.57 | 3.18 | 3.48 | 0.50 | 11.50 | S. epidermidis 2.6 |
| 53.96 | 21.01 | 2.06 | 3.23 | 5.25 | 0.45 | 14.00 | S. epidermidis 2.6 |
| 7.02 | 2.67 | 0.28 | 0.48 | 0.21 | 0.07 | 89.30 | S. epidermidis mix |
| 6.48 | 2.66 | 0.27 | 0.45 | 0.22 | 0.06 | 89.90 | S. epidermidis mix |
| 7.94 | 2.37 | 0.27 | 0.46 | 0.13 | 0.07 | 88.80 | S. epidermidis mix |
| 41.34 | 15.68 | 1.36 | 2.29 | 31.51 | 0.43 | 7.40 | S. epidermidis CA7 |
| 49.98 | 15.60 | 1.47 | 2.41 | 22.00 | 0.44 | 8.10 | S. epidermidis CA7 |
| 50.73 | 15.34 | 1.60 | 1.99 | 23.47 | 0.46 | 6.41 | S. epidermidis CA7 |
| 68.13 | 9.92 | 1.92 | 2.84 | 0.00 | 0.80 | 16.40 | TSB control |
| 64.30 | 10.55 | 2.35 | 2.75 | 0.00 | 0.79 | 19.30 | TSB control |
| 74.81 | 8.87 | 2.14 | 2.88 | 0.00 | 0.79 | 10.50 | TSB control |

Note: IA: Indole-3-acrylic acid; IAA: Indole-3-acetic acid; Iald: Indole-3-aldehyde; IE: Indole-3-ethanol; ILA: Indole-3-lactic acid; IPA: Indole-3-propionic acid; TRYA: Tryptamine.

**Table S5.** Reproducibility test of standard with five replicates for the 4-min method. The method showed an average CV% of 5 % for quantification and an average CV% of 0.3% for retention time, indicating the robustness of the high-throughput method.

| Injection volume (µL) | Average CV% for quantification | Average CV% for retention time |
|---|---|---|
| 0.2 | 6.44 | 0.32 |
| 0.5 | 5.00 | 0.26 |
| 1.0 | 4.60 | 0.26 |
| 25.0 | 5.56 | 0.32 |

**Table S6.** Quantified indole derivatives of 24 *Bacteroides uniformis* strains.

| IAM | IAA | IAcA | ICA | IEH | TRM | Label |
|---|---|---|---|---|---|---|
| 1.33 | 19.21 | 28.53 | 37.89 | 7.75 | 5.29 | ME/CFS |
| 1.37 | 16.76 | 18.73 | 47.29 | 8.48 | 7.39 | Control |
| 0.89 | 19.1 | 44.47 | 23.69 | 5.35 | 6.5 | ME/CFS |
| 1.51 | 17.5 | 26.4 | 39.92 | 8.86 | 5.81 | ME/CFS |
| 1.04 | 20.6 | 29.23 | 35.14 | 7.57 | 6.43 | Control |
| 0.89 | 21.22 | 38.4 | 26.64 | 6.37 | 6.48 | Control |
| 0.95 | 20.66 | 32.51 | 34.18 | 6.02 | 5.67 | Control |
| 1.63 | 12.57 | 10.84 | 58.37 | 7.59 | 9 | ME/CFS |
| 0.99 | 23.7 | 27.42 | 32.84 | 8.29 | 6.76 | ME/CFS |
| 0.76 | 25.89 | 29.3 | 30.52 | 6.81 | 6.73 | ME/CFS |
| 1.24 | 24.82 | 22.31 | 35.63 | 8.53 | 7.47 | ME/CFS |
| 1.09 | 22.08 | 25.06 | 37.1 | 7.07 | 7.6 | Control |
| 0.94 | 21.21 | 35.6 | 28.67 | 7.08 | 6.5 | Control |
| 1.18 | 23.78 | 17.59 | 40.01 | 8.16 | 9.28 | ME/CFS |
| 0.93 | 26.43 | 23.95 | 35.43 | 6.45 | 6.81 | Control |
| 0.57 | 21.38 | 48.26 | 19.22 | 4.31 | 6.26 | Control |
| 0.88 | 22.91 | 35.34 | 27.78 | 5.54 | 7.56 | ME/CFS |
| 1.37 | 20.11 | 11.21 | 50.34 | 9.83 | 7.15 | ME/CFS |
| 1.14 | 16.69 | 27.42 | 39.83 | 7.01 | 7.91 | Control |
| 1.22 | 18.11 | 24.43 | 41.5 | 7.92 | 6.81 | Control |
| 0.61 | 22.59 | 44.75 | 22.58 | 4.49 | 4.99 | ME/CFS |
| 0.85 | 16.19 | 41.68 | 30.37 | 5.97 | 4.92 | Control |
| 0.94 | 24.75 | 38.8 | 23.33 | 5.56 | 6.61 | ME/CFS |
| 0.44 | 18.28 | 55.81 | 17.06 | 3.69 | 4.72 | Control |

Note: IAcA: Indole-3-acrylic acid; IAA: Indole-3-acetic acid; ICA: Indole-3-aldehyde; IEH: Indole-3-ethanol; TRM: Tryptamine.

**Table S7.** Quantified compounds of the bacterial indole pathway in fecal samples.

| IEH | IPA | IAA | ICA | TRM | Label |
|---:|---:|---:|---:|---:|---:|
| 11.81 | 45.19 | 19.06 | 19.39 | 4.54 | Control |
| 4.97 | 12.48 | 27.52 | 5.67 | 49.37 | Control |
| 1.71 | 21.69 | 25.49 | 1.97 | 49.14 | Control |
| 3.23 | 66.96 | 20.26 | 6.23 | 3.32 | Control |
| 13.22 | 33.53 | 31.33 | 19.13 | 2.79 | Control |
| 7.09 | 39.49 | 21.01 | 25.46 | 6.94 | Control |
| 3.65 | 29.99 | 19.52 | 45.13 | 1.70 | Control |
| 2.43 | 4.05 | 7.44 | 2.89 | 83.19 | Control |
| 12.12 | 28.62 | 26.95 | 29.86 | 2.45 | Control |
| 12.42 | 16.63 | 57.69 | 8.42 | 4.83 | Control |
| 0.75 | 23.48 | 71.41 | 2.36 | 1.99 | ME/CFS |
| 2.25 | 20.03 | 71.61 | 3.42 | 2.69 | ME/CFS |
| 8.46 | 18.82 | 31.75 | 5.04 | 35.93 | ME/CFS |
| 8.69 | 55.09 | 23.76 | 8.97 | 3.48 | ME/CFS |
| 8.10 | 40.66 | 25.41 | 0.81 | 25.02 | ME/CFS |
| 0.47 | 41.80 | 16.83 | 40.80 | 0.10 | ME/CFS |
| 5.97 | 49.34 | 10.69 | 14.46 | 19.54 | ME/CFS |
| 6.96 | 23.49 | 34.63 | 29.68 | 5.24 | ME/CFS |
| 1.30 | 9.53 | 17.57 | 1.64 | 69.96 | ME/CFS |
| 10.90 | 23.17 | 23.24 | 38.74 | 3.96 | ME/CFS |

Note: IAA: Indole-3-acetic acid; ICA: Indole-3-aldehyde; IEH: Indole-3-ethanol; IPA: Indole-3-propionic acid; TRM: Tryptamine.

**Figure S1**. Calibration curves for the 28-min method were achieved for different indole compounds by plotting the intensity ratio of each compound over NFK-C13 vs. the concentration ratio. The $R^2$ values were shown in Table 1. Each concentration was measured in triplicates. DIC: 5,11-dihydroindolo[3,2-b]carbazole; IAA: Indole-3-acetic acid; IAcA: Indole-3-acrylic acid; IAM: Indole-3-acetamide; IAN: Indole-3-acetonitrile; IBA: indole-3-butyric acid; ICA: Indole-3-aldehyde; IEH: Indole-3-ethanol; ILA: Indole-3-lactic acid; IND: Indole; IPA: Indole-3-propionic acid; KYN: Kynurenine; NFK: N-Formylkynurenine; TRM: Tryptamine.

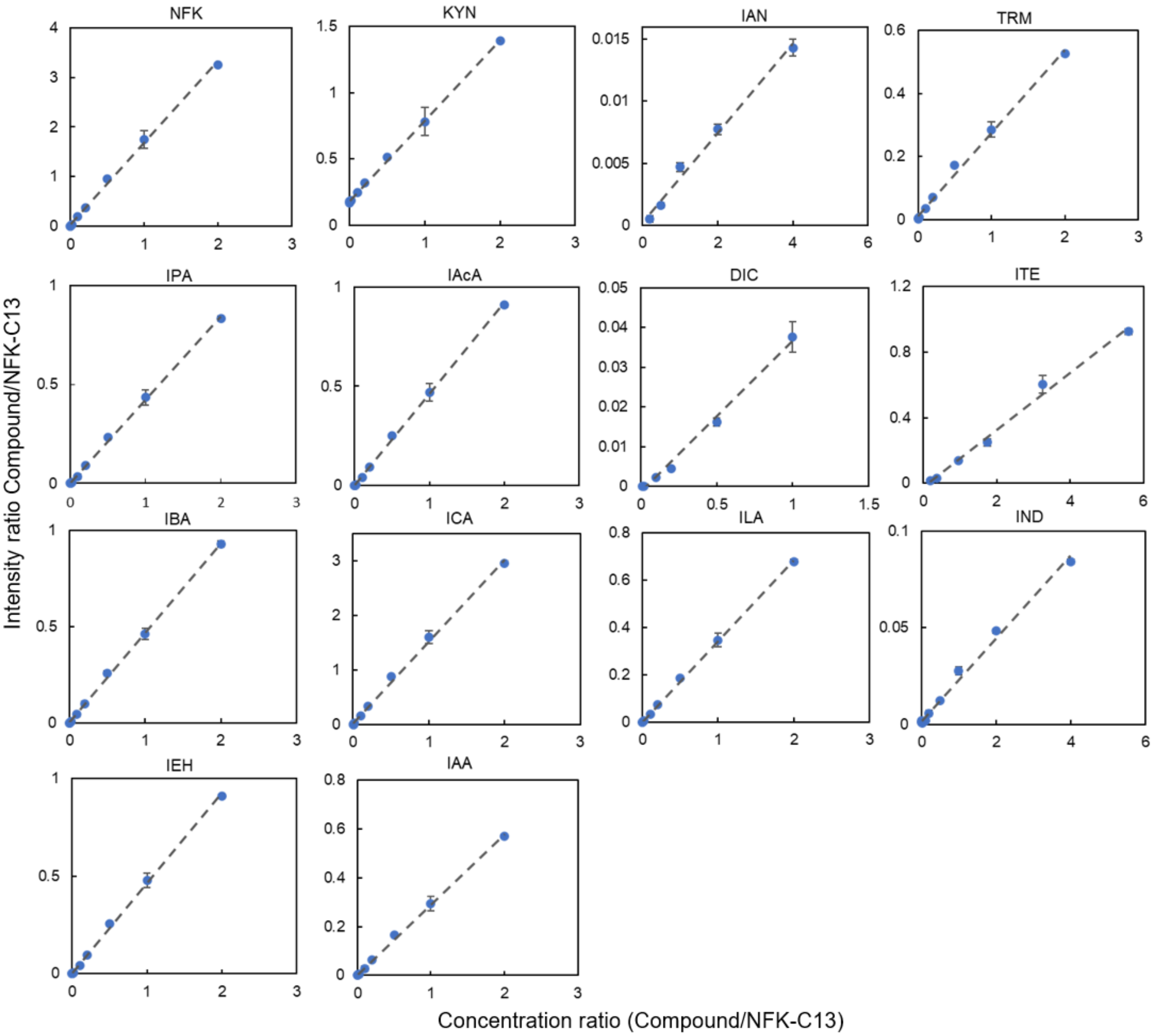

**Figure S2.** Structure and isotope overlap of light and heavy tryptophan catabolites. (A) N-Formylkynurenine (NFK and NFK-C13). (B) Indole-3-carboxaldehyde (ICA and ICA-8C13). (C) Indole-3-pronoic acid (IPA and IPA-2D).

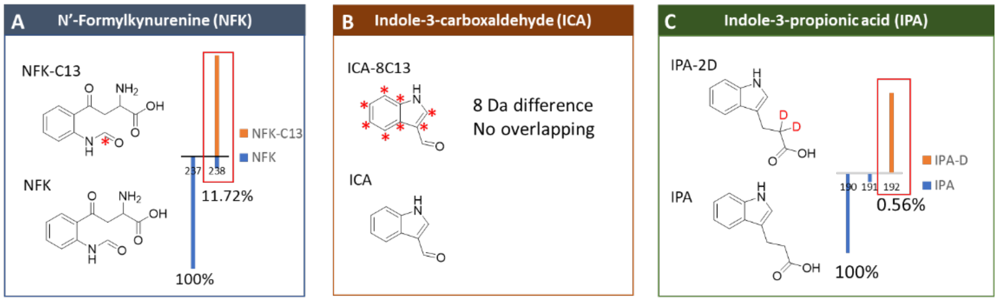

**Figure S3**. Calibration curves for kynurenine using different internal standards. (A) N-Formylkynurenine-C13 (NFK-C13). (B) Indole-3-carboxaldehyde-8C13 (ICA-8C13). (C) Indole-3-pronoic acid-2D (IPA-2D).

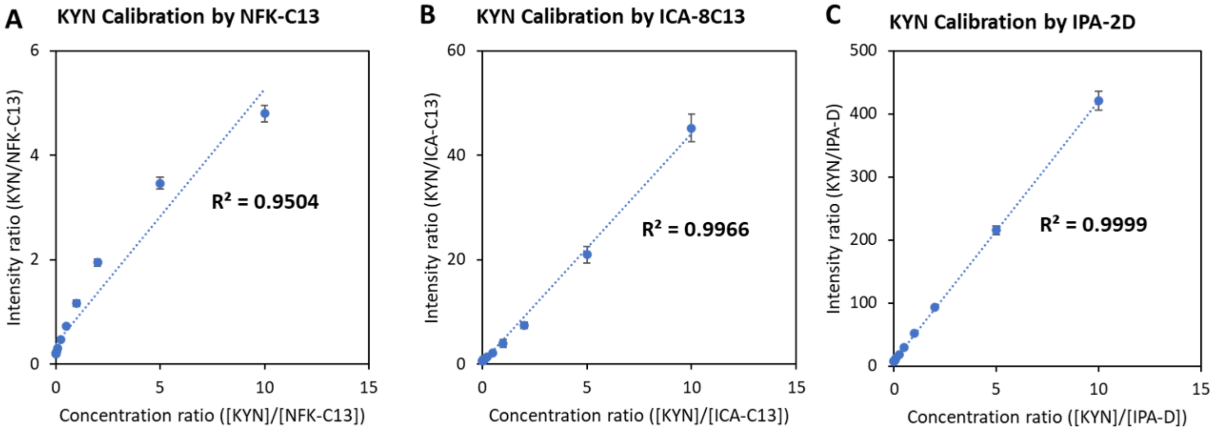

**Figure S4**. Calibration curves for the 4-min method were achieved for different indole compounds by plotting the intensity ratio of each compound over ICA-8C13 vs. the concentration ratio. The R2 values were shown in Table 1. Each concentration was measured in triplicates. 3-HAA: 3-Hydroxynthranileic acid; IAA: Indole-3-acetic acid; IAcA: Indole-3-acrylic acid; IAM: Indole-3-acetamide; IAN: Indole-3-acetonitrile; IBA: indole-3-butyric acid; ICA: Indole-3-aldehyde; IEH: Indole-3-ethanol; ILA: Indole-3-lactic acid; IPA: Indole-3-propionic acid; KYN: Kynurenine; NFK: N-Formylkynurenine; TRM: Tryptamine.

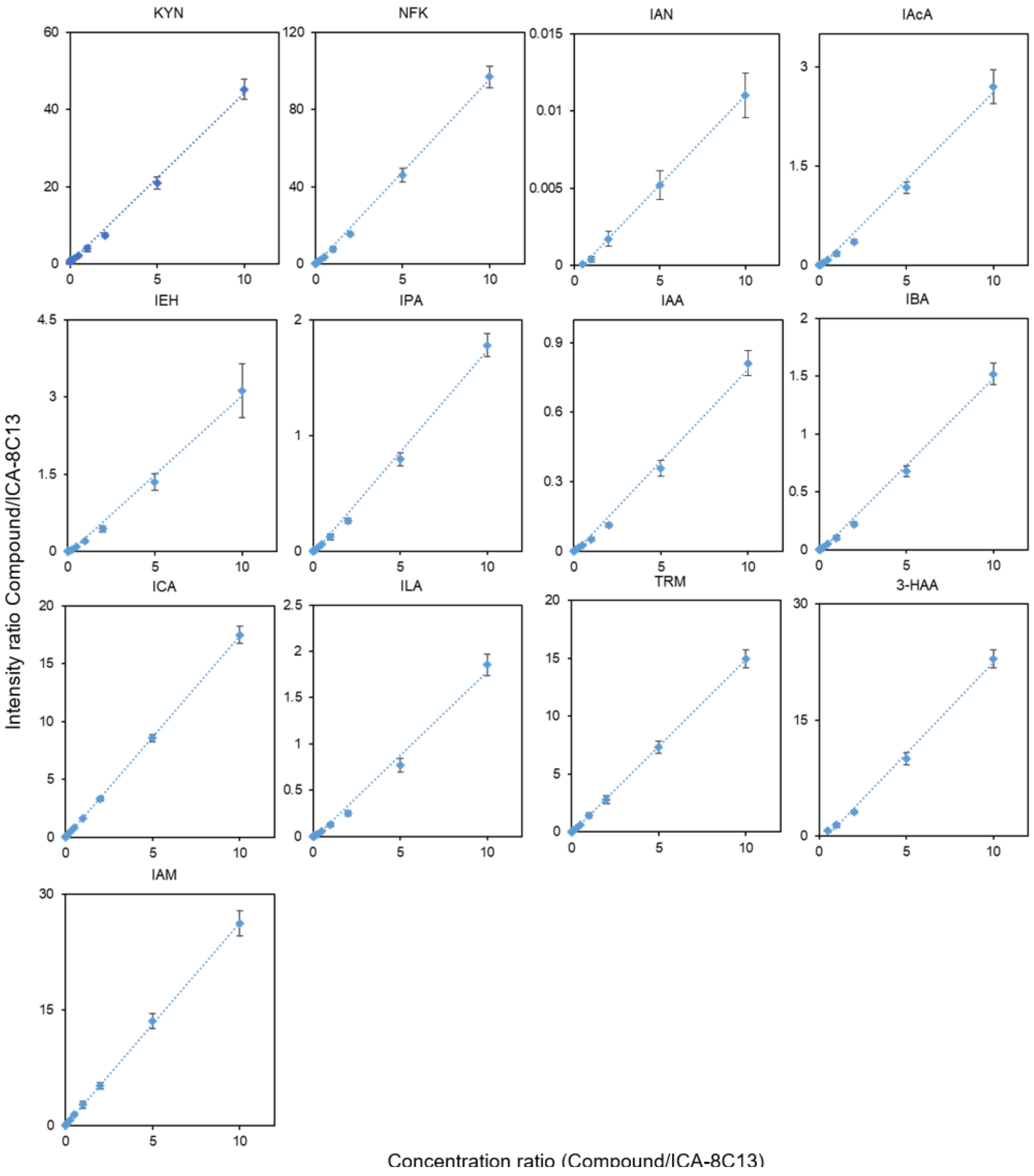

**Figure S5**. Screenshots of ion chromatogram and skyline ion chromatogram of the old 28-min method and the new 4-min method. Both methods had satisfactory resolution. A similar separation was achieved by the high-throughput method with a clean ion chromatogram (15k in resolution at 200 *m/z* for the 28-min method and 20k for the 4-min method).

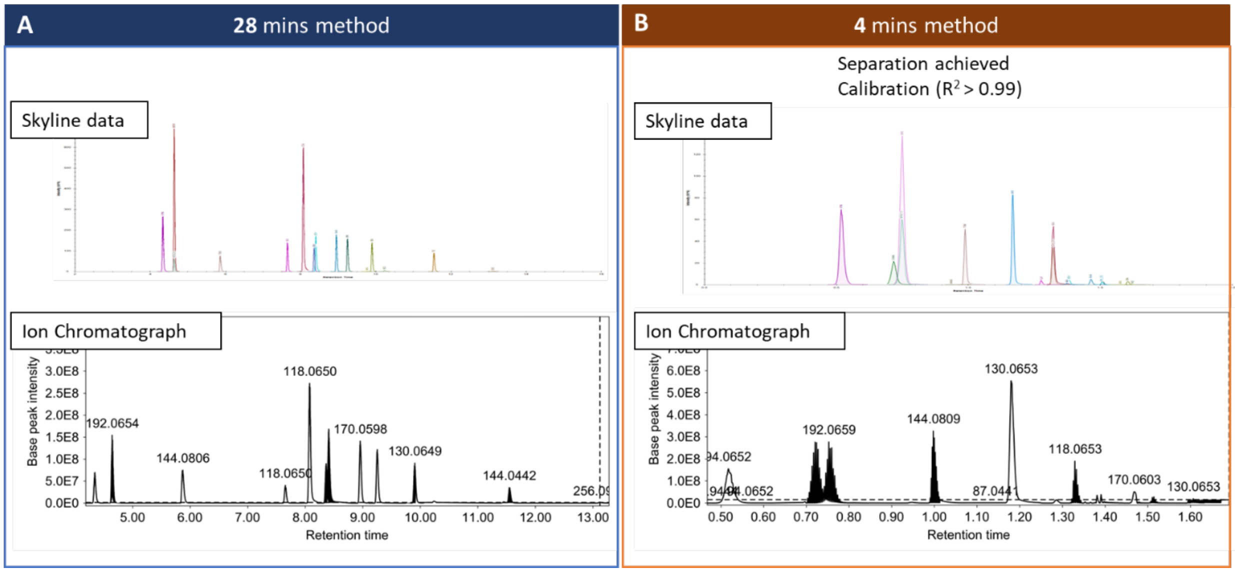

**Scheme S1**. Bacterial tryptophan degradation. (Lu, Y.; Chong, J.; Shen, S.; Chammas, J.-B.; Chalifour, L.; Xia, J. TrpNet: Understanding Tryptophan Metabolism across Gut Microbiome. Metabolites 2021, 12 (1), 10. https://doi.org/10.3390/metabo12010010.)

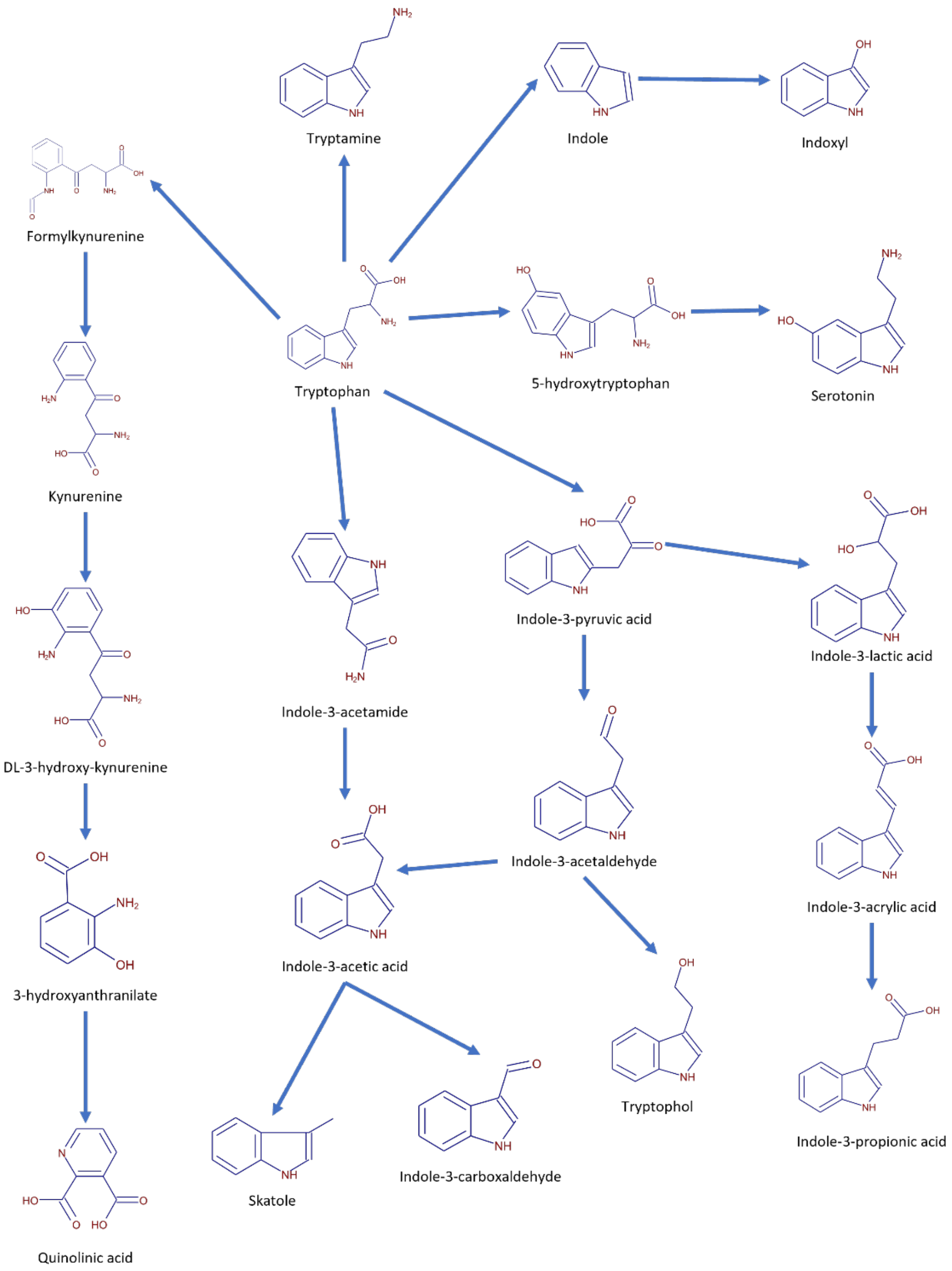

**Scheme S2**. Superpathway of indole-3-acetate conjugate biosynthesis.

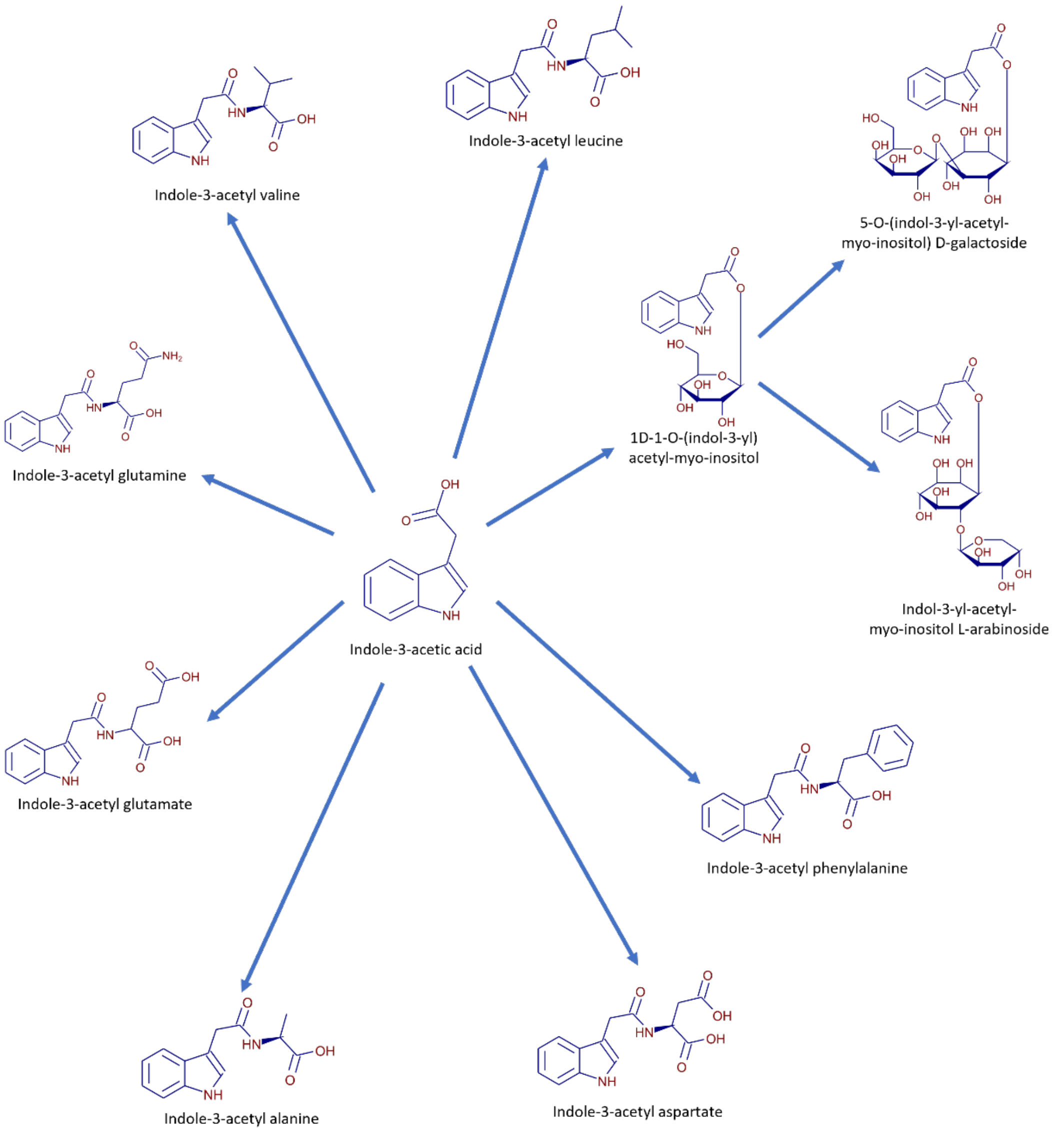
